# pUL21 regulation of pUs3 Kinase Activity Influences the Nature of Nuclear Envelope Deformation by the HSV-2 Nuclear Egress Complex

**DOI:** 10.1101/2021.06.02.446771

**Authors:** Jamil H. Muradov, Renée L. Finnen, Michael A. Gulak, Thomas J. M. Hay, Bruce W. Banfield

## Abstract

It is well established that the herpesvirus nuclear egress complex (NEC) has an intrinsic ability to deform membranes. During viral infection, the membrane-deformation activity of the NEC must be precisely regulated to ensure efficient nuclear egress of capsids. One viral protein known to regulate herpes simplex virus type 2 (HSV-2) NEC activity is the tegument protein pUL21. Cells infected with an HSV-2 mutant lacking pUL21 (ΔUL21) produced a slower migrating species of the viral serine/threonine kinase pUs3 that was shown to be a hyperphosphorylated form of the enzyme. Investigation of the pUs3 substrate profile in ΔUL21-infected cells revealed a prominent band with a molecular weight consistent with that of the NEC components pUL31 and pUL34. Phosphatase sensitivity and retarded mobility in phos-tag SDS-PAGE confirmed that both pUL31 and pUL34 were hyperphosphorylated by pUs3 in the absence of pUL21. To gain insight into the consequences of increased phosphorylation of NEC components, the architecture of the nuclear envelope in cells producing the HSV-2 NEC in the presence or absence of pUs3 was examined. In cells with robust NEC production, invaginations of the inner nuclear membrane were observed that contained budded vesicles of uniform size. By contrast, nuclear envelope deformations protruding outwards from the nucleus, were observed when pUs3 was included in transfections with the HSV-2 NEC. Finally, when pUL21 was included in transfections with the HSV-2 NEC and pUs3, decreased phosphorylation of NEC components was observed in comparison to transfections lacking pUL21. These results demonstrate that pUL21 influences the phosphorylation status of pUs3 and the HSV-2 NEC and that this has consequences for the architecture of the nuclear envelope.

**Author Summary:** During all herpesvirus infections, the nuclear envelope undergoes deformation in order to enable viral capsids assembled within the nucleus of the infected cell to gain access to the cytoplasm for further maturation and spread to neighbouring cells. These nuclear envelope deformations are orchestrated by the viral nuclear egress complex (NEC), which, in HSV, is composed of two viral proteins, pUL31 and pUL34. How the membrane-deformation activity of the NEC is controlled during infection is incompletely understood. The studies in this communication reveal that the phosphorylation status of pUL31 and pUL34 can determine the nature of nuclear envelope deformations and that the viral protein pUL21 can modulate the phosphorylation status of both NEC components. These findings provide an explanation for why HSV-2 strains lacking pUL21 are defective in nuclear egress. A thorough understanding of how NEC activity is controlled during infection may yield strategies to disrupt this fundamental step in the herpesvirus lifecycle, providing the basis for novel antiviral strategies.

## Introduction

The assembly pathway of all herpesvirus virions begins inside the nucleus of infected cells. Newly synthesized viral genomes are packaged into preformed procapsids and then DNA-containing capsids, called C-capsids, are preferentially selected for subsequent incorporation into mature virions. C-capsids must transit from the nucleoplasm across the nuclear envelope (NE) and be deposited in the cytoplasm where the final stages of virion maturation take place. C-capsids acquire a primary envelope at the inner nuclear membrane (INM) resulting in the formation of primary enveloped virions (PEVs) within the perinuclear space. PEVs then fuse with the outer nuclear membrane (ONM) releasing the C-capsids into the cytoplasm. This process, referred to as nuclear egress, is facilitated by the highly conserved nuclear egress complex (NEC). In herpes simplex virus (HSV) the NEC is comprised of two viral proteins, pUL31 and pUL34. Mutant HSV-1 strains deleted for UL31 or UL34 display a 4-log reduction in virus replication in most cell types and an accumulation of capsids in the nucleus [1, 2].

pUL34 is a type II transmembrane protein with a C-terminal domain anchored to the nucleoplasmic face of the INM as well as the cytoplasmic face of the endoplasmic reticulum (ER). pUL31 is a soluble nuclear phosphoprotein that can be recruited to the INM through its interaction with the N-terminus of pUL34. In pseudorabies virus (PRV) and HSV-1, pUL31 can be co-purified with capsids and it has been suggested that capsid-associated pUL31 can mediate the recruitment of C-capsids to the INM via its interactions with pUL34 [3–5]. Structural studies of the NEC revealed that, once bound to each other at the INM, pUL31 and pUL34 form hexamers of NEC heterodimers which arrange into a honeycomb lattice that can introduce negative curvature of the membrane [6, 7]. The capacity of the NEC to deform membranes is crucial for capsid envelopment at the INM. In cells transfected with PRV pUL31 and pUL34 expression plasmids, the NEC is active and vesicles derived from the INM accumulate within herniations of the perinuclear space [8]. By contrast, in infected cells pUL31 and pUL34 co-localize predominantly at the INM and vesiculation of the INM is not apparent [9]. These findings indicate that the membrane deformation activity of the NEC is regulated by other proteins in virally infected cells to prevent excessive vesiculation of the INM. It is likely that the NEC lattice at the INM remains quiescent and preserves the integrity of the INM until such time as a C-capsid arrives at the site of primary envelopment, activating the NEC [10].

A promiscuous viral serine/threonine kinase, called pUs3, is implicated in regulation of multiple host-virus interactions, but its most well-known role is in nuclear egress [11]. In the absence of pUs3 activity, many alphaherpesviruses accumulate PEVs in the perinuclear space within herniations of the INM [12–15]. These findings have largely been interpreted to suggest a role for pUs3 in the de-envelopment of PEVs at the ONM. A recent study suggested that pUs3 drives the formation of INM herniations that serve as sites for primary envelopment of capsids, however, the resulting PEVs are too far away from the ONM to allow for de-envelopment [16]. pUs3 phosphorylates both pUL31 and pUL34 [17, 18]. While the role of pUL34 phosphorylation in nuclear egress is unclear, pUs3 phosphorylation of serine residues in the N-terminus of HSV-1 pUL31 regulates NEC activity [19, 20]. Similar to what is observed in pUs3 mutant viruses, substitution of the N-terminal pUL31 serine residues for alanine resulted in the accumulation of PEVs in INM herniations whereas phosphomimetic substitution of these residues for glutamic acid interfered with primary envelopment of capsids [19].

Recent work from this laboratory suggested that pUL21 also regulates the NEC. In multiple HSV-2 strains, mutations in UL21 resulted in 100-fold or greater replication deficiencies and defects in the nuclear egress of capsids [16, 21, 22]. By contrast, HSV-1 strains deleted for UL21 show more modest reductions in virus replication [16, 21, 23, 24]. Notably, distribution of the NEC and the architecture of the NE was altered in both HSV-1 and HSV-2 UL21 mutants [16]. Importantly, knock-down of pUL31 production in cells infected with HSV-2 UL21 mutant viruses prevented NE deformations indicating that pUL31 activity was required for the formation of these structures and suggesting that pUL21 can regulate NEC activity [16].

The goals of the present study were to understand how pUL21 regulates the HSV-2 NEC. We found that in cells infected with an HSV-2 UL21 mutant, not only pUs3 but also its substrates, pUL31 and pUL34, were hyperphosphorylated. Investigation into the consequences of pUL31 and pUL34 phosphorylation on NEC activity indicated that hypophosphorylation of NEC components was associated with invagination of the INM and INM vesiculation, whereas phosphorylation of NEC components resulted in extravagation of NE components into the cytoplasm. This study provides insight into the mechanism by which pUL21 and pUs3 regulate NEC activity and the impact of phosphorylation of NEC components on NE architecture.

## Materials and Methods

### Cells and viruses

African green monkey kidney cells (Vero), HeLa cells, human keratinocytes (HaCaT) and HaCaT21 cells, which stably express HSV-2 186 pUL21 [16], were maintained in Dulbecco’s modified Eagle medium (DMEM) supplemented with 10% fetal bovine serum (FBS) in a 5% CO_2_ environment. HSV-2 strain 186 was acquired from Dr. Yasushi Kawaguchi (University of Tokyo). HSV-2 strain 186 viruses deficient in pUL21 (ΔUL21) and pUs3 (ΔUs3) were constructed as described previously [16, 22]. An HSV-2 strain 186 virus deficient in both pUL21 and pUs3 (ΔUL21/ΔUs3) was constructed by two-step Red-mediated mutagenesis of a Us3 null BAC using primers to introduce a deletion into the UL21 gene as described previously [22]. HSV-2 strain 186 viruses repaired for deficiency in pUL21 (ΔUL21R) or pUs3 (ΔUs3R) were constructed as described previously [16, 22]. All pUL21 deficient viruses were propagated in HaCaT21 cells. Times post infection reported as hours post infection refer to the time elapsed following medium replacement after a one-hour inoculation period.

### Plasmids

The construction of plasmids encoding wild type pUs3 and a D305A mutant of pUs3 lacking kinase activity (Us3KD) was described previously [25]. The construction of a plasmid encoding EGFP-pUL31 was described previously [26]. To change serines 9, 11, 24, 27, 39, 42 and 45 and threonine 20 of EGFP-pUL31 to alanine, the primers listed in Table 1 were used in successive mutagenic PCR reactions. The construction of a plasmid encoding Flag-pUL34 was described previously [26]. To construct a plasmid encoding untagged pUL34, PCR utilizing forward primer 5’-GATCGAATTCATGGCGGGGATGGGGAAGC-3’ and reverse primer 5’-GATCGTCGACTCATAGGCGCGCGCCAAC-3’ were used to amplify the UL34 gene using purified HSV-2 DNA as template. The product was digested with EcoRI and SalI and ligated into similarly digested pCI-neo (Promega, Madison, WI). To change serines 195 and 198 of pUL34 to alanine, the primers listed in Table 1 were used in successive mutagenic PCR reactions. The construction of a plasmid encoding pUL21 was described previously [22]. To construct a plasmid encoding an mCherry tagged lamin associated protein 2β (mCh-LAP2β), PCR utilizing forward primer 5’-GATCACCGGTATGGTGAGCAAGGGCGAGG-3’ and reverse primer 5’-GATCGTCGACCTTGTACAGCTCGTCCATGCCCG-3’ were used to amplify the mCherry gene using plasmid pJR70 [25] as template. The product was digested with AgeI and SalI and ligated into similarly digested plasmid encoding EGFP-LAP2β, generously provided by Dr. Tokuko Haraguchi (Osaka University). All plasmids constructed utilizing PCR were sequenced to ensure that no unintended mutations were introduced.

**Table 1.**
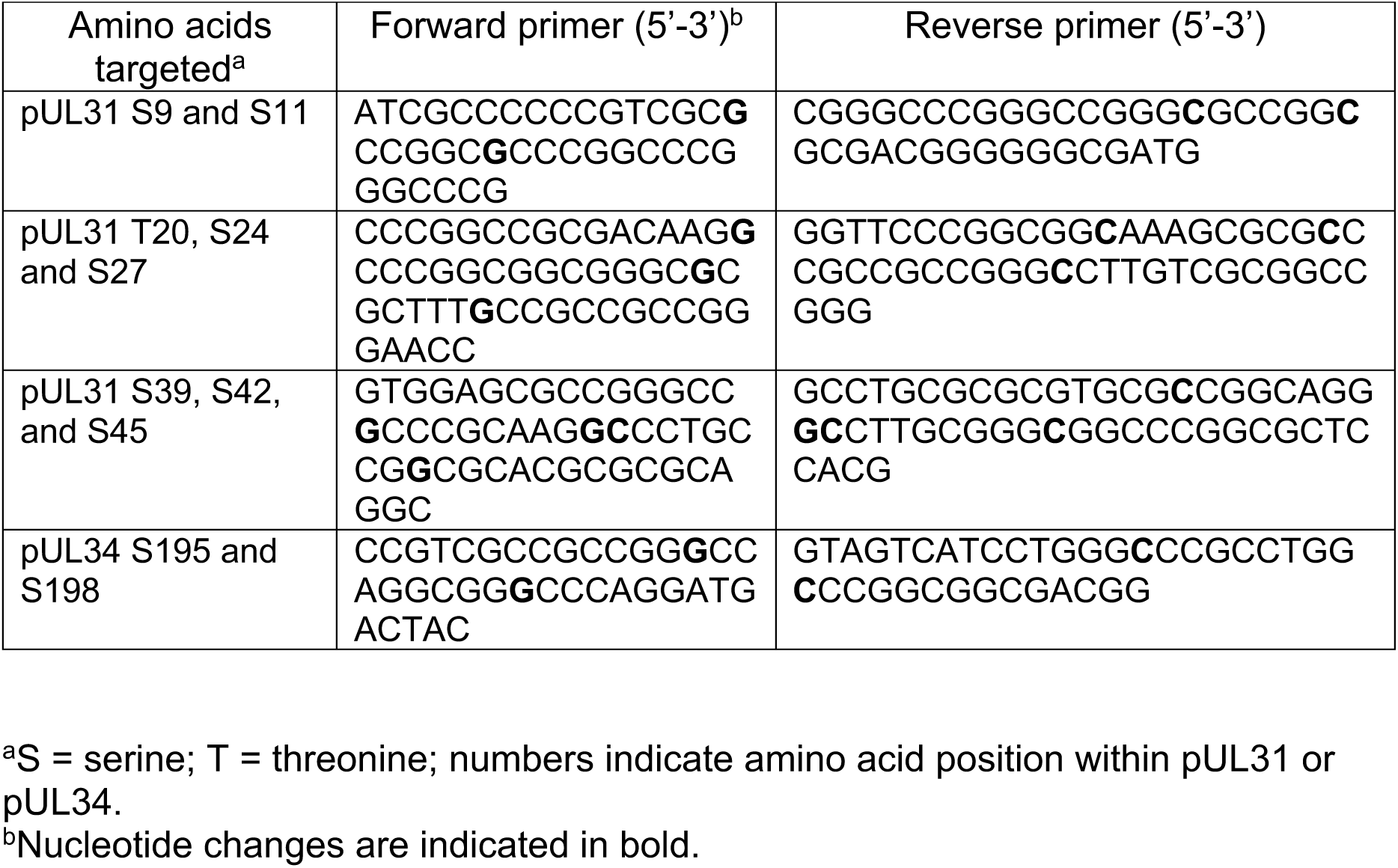
Oligonucleotide primers used for mutagenesis of EGFP-pUL31 and pUL34

### Immunological reagents

Rat polyclonal antiserum against pUL31 [16] was used for western blotting at a dilution of 1:200; chicken polyclonal antiserum against pUL34 [16] was used for western blotting at a dilution of 1:500; rat polyclonal antiserum against pUs3 [25] was used for western blotting at a dilution of 1:1,000; rat polyclonal antiserum against pUL21 [22] was used for western blotting at a dilution of 1:600; mouse monoclonal antibody against ICP27 (Virusys, Taneytown, MD) was used for western blotting at a dilution of 1:500, mouse monoclonal antibody against β-actin (Sigma, St. Louis, MO) was used for western blotting at a dilution of 1:2,000; mouse monoclonal antibody against EGFP (Clontech, Mountain View, CA) was used for western blotting at a dilution of 1:1,000 and rabbit polyclonal antiserum reactive against protein kinase A (PKA) substrates (Cell Signaling Technology, Danvers, MA) was used for western blotting at a dilution of 1:1,000. Horseradish peroxidase-conjugated goat anti-chicken IgY and horseradish peroxidase-conjugated rabbit anti-rat IgG (Sigma, St. Louis, MO) were used for western blotting at a dilution of 1:80,000. Horseradish peroxidase-conjugated goat anti-rabbit IgG (Sigma, St. Louis, MO) was used for western blotting at a dilution of 1:10,000. Horseradish peroxidase-conjugated goat anti-mouse IgG (Jackson ImmunoResearch, West Grove, PA) was used for western blotting at a dilution of 1:90,000. Mouse monoclonal antibody against lamin A/C (EMD, Millipore Temecula, CA) was used for indirect immunofluorescence microscopy at a dilution of 1:300 and mouse monoclonal antibody against nuclear pore complex proteins (Abcam, Cambridge, MA) was used for indirect immunofluorescence microscopy at a dilution of 1:500. Alexa Fluor 568 conjugated donkey anti-mouse IgG (Molecular Probes, Eugene, OR) was used for indirect immunofluorescence microscopy at a dilution of 1:500.

### Preparation and analysis of cellular protein extracts

Whole cell extracts of infected cells were prepared by rinsing cell monolayers with cold phosphate buffered saline (PBS), followed by scraping into cold PBS containing protease inhibitors (Roche, Laval, QC). Harvested cells were transferred to 1.5 ml tubes containing 3X SDS-PAGE loading buffer. The lysate was repeatedly passed through a 28 ½-gauge needle to reduce viscosity and then heated at 100°C for 5 minutes. For western blot analysis, 10 to 20 μl of whole cell extract were subjected to SDS-PAGE. Separated proteins were transferred to PVDF membranes (Millipore, Billerica, MA) and probed with appropriate dilutions of primary antibody followed by appropriate dilutions of horseradish peroxidase-conjugated secondary antibody. The membranes were treated with Pierce ECL western blotting substrate (Thermo Scientific, Rockford, IL) and exposed to film.

To prepare cellular extracts for analyzing proteins by phos-tag SDS-PAGE, infected or transfected monolayers of cells grown in 35 mm dishes were washed with cold PBS and then scraped into 200 μl cold RIPA buffer (50 mM Tris-HCl pH 7.5, 150 mM NaCl, 0.5% sodium deoxycholate, 1% NP-40, 0.1% SDS) containing protease inhibitors (Roche, Laval, QC). The lysates were repeatedly passed through 28 ½-gauge needles to reduce viscosity and 250 units of benzonase nuclease (Santa Cruz Biotechnology, Dallas, TX) were added to each lysate. The lysates were centrifuged for 10 minutes at 14,000 x g and the supernatants were transferred to 1.5 ml tubes containing 100 μl 3X SDS-PAGE loading buffer. For lambda protein phosphatase (λ PP) treatment of lysates, 39 μl of each lysate was combined with 5 μl of 10X NEB buffer for Protein MetalloPhosphatases (New England Biolabs, Ipswich, MA), 5 μl of 10 mM MnCl_2_ and 1-2 μl (400-800 units) of λ PP (New England Biolabs, Ipswich, MA). Untreated lysates received 1-2 μl of water in place of λ PP. The dephosphorylation reactions were carried out for 30 to 45 minutes at 30°C. For western blot analysis, 10 to 20 μl of treated lysates were electrophoresed through gels containing 50 μM Phos-tag reagent (Fujifilm, Richmond, USA) and 100 µM MnCl_2_. After electrophoresis, gels were soaked in transfer buffer containing 1 mM EDTA, re-equilibrated in transfer buffer and proteins were transferred to PVDF membranes and probed as described above.

### Fluorescence and immunofluorescence microscopy of transfected cells

To examine the localization of EGFP-pUL31 by microscopy, HeLa cells growing on 35 mm glass bottom dishes (MatTek, Ashland, MA) were transfected using X-treme GENE HP DNA transfection reagent (Roche, Laval, QC) according to manufacturer’s instructions. At 18 hours post transfection, cells were fixed with 4% paraformaldehyde in PBS and stained as described previously (Finnen et al., 2010). Images were captured with an Olympus FV1000 laser scanning confocal microscope using a 60X (1.42 NA) oil immersion objective lens and Fluoview 1.7.3.0 software. Composites of representative images were prepared using Adobe Photoshop software. To quantitate extravagation formation in transfected cells, multiple images of fixed cells stained with Hoechst 33342 (Sigma, St. Louis, MO) were captured using a fixed PMT voltage for the EGFP channel. Using Image Pro software, the level of fluorescence in an area of interest (AOI) encompassing all extravagations (AOI1) was measured. The perimeter of the nucleus was defined based on the Hoechst 33342 channel and then the level of fluorescence inside the nucleus (AOI2) was measured. The level of fluorescence in AOI2 was subtracted from the level of fluorescence in AOI1 to provide a measure of the level of fluorescence outside the nucleus that was expressed as a percentage of the level of fluorescence in AOI1.

### Fluorescence recovery after photobleaching analyses

HeLa cells growing on 35 mm glass bottom dishes (MatTek, Ashland, MA) were transfected using X-treme GENE HP DNA transfection reagent (Roche, Laval, QC) according to manufacturer’s protocols. At 18 hours post transfection, cells were mounted onto an Olympus FV1000 laser scanning confocal microscope and maintained at 37°C in a humidified 5% CO_2_ environment. All image acquisition parameters were controlled using Fluoview software version 1.7.3.0. EGFP was excited using a 488 ηm laser set at 5% power. Images at 512-pixel-by-512-pixel resolution were collected using a 60X (1.42 NA) oil immersion objective lens at a rate of 1.1 frames per second. Frames were collected before the indicated region was photobleached by repeated scanning of the region of interest with a 405 ηm laser set at 100% power for a total of 300 ms. After photobleaching, additional frames were collected for 5 minutes. The fluorescence intensity in bleached and unbleached control regions in each frame were measured using Fluoview software version 1.7.3.0. and the data were exported into Microsoft Excel for graphical presentation.

### Transmission electron microscopy

Plasmids were transfected into HeLa cells growing on 100 mm dishes using X-treme GENE HP DNA transfection reagent (Roche, Laval, QC) according to manufacturer’s protocols and processed for TEM at 18 hours post transfection. Transfected cells were washed with PBS three times before fixing in 1.5 ml of 2.5% EM grade glutaraldehyde (Ted Pella, Redding, CA) in 0.1 M sodium cacodylate buffer (pH 7.4) for 60 minutes. Cells were collected by scraping into fixative and centrifugation at 300 x g for five minutes. Cell pellets were carefully enrobed in an equal volume of molten 5% low-melting temperature agarose (Lonza, Rockland, ME) and allowed to cool. Specimens in agarose were incubated in 2.5% glutaraldehyde in 0.1 M sodium cacodylate buffer (pH 7.4) for 1.5 hours and post-fixed in 1% osmium tetroxide for one hour. The fixed cells in agarose were washed with distilled water three times and stained in 0.5% uranyl acetate overnight before dehydration in ascending grades of ethanol (30%-100%). Samples were transitioned from ethanol to infiltration with propylene oxide and embedded in Embed-812 hard resin (Electron Microscopy Sciences, Hatfield, PA). Blocks were sectioned at 50-60 ηm and stained with uranyl acetate and Reynolds’ lead citrate. Images were collected using a Hitachi H-7000 transmission electron microscope.

## Results

### Deletion of pUL21 from HSV-2 results in pUs3 phosphorylation and an increase in pUs3-mediated phosphorylation of NEC components

During our characterization of an HSV-2 strain 186 virus lacking pUL21 (ΔUL21), we noted that a slower migrating species of pUs3, indicated with an asterisk in Figure 1A, was present in cells infected with ΔUL21 but not in cells infected with parental 186 virus. As phosphorylated forms of pUs3 from other alphaherpesviruses have been described previously [27, 28], we speculated that this slower migrating species was phosphorylated HSV-2 pUs3. Phosphatase treatment of lysates prior to western blotting resulted in the disappearance of the slower migrating pUs3 species indicating that pUs3 is more heavily phosphorylated in the absence of pUL21 (Figure 1B). To corroborate these findings, cell lysates were electrophoresed through phos-tag gels prior to western blotting (Figure 1C). In ΔUL21 infected cell lysates a distinct, slower migrating, phosphatase sensitive pUs3 species was detected that was largely absent in 186 infected cell lysates. These findings confirm that pUs3 is hyperphosphorylated in the absence of pUL21.

**Figure 1.**
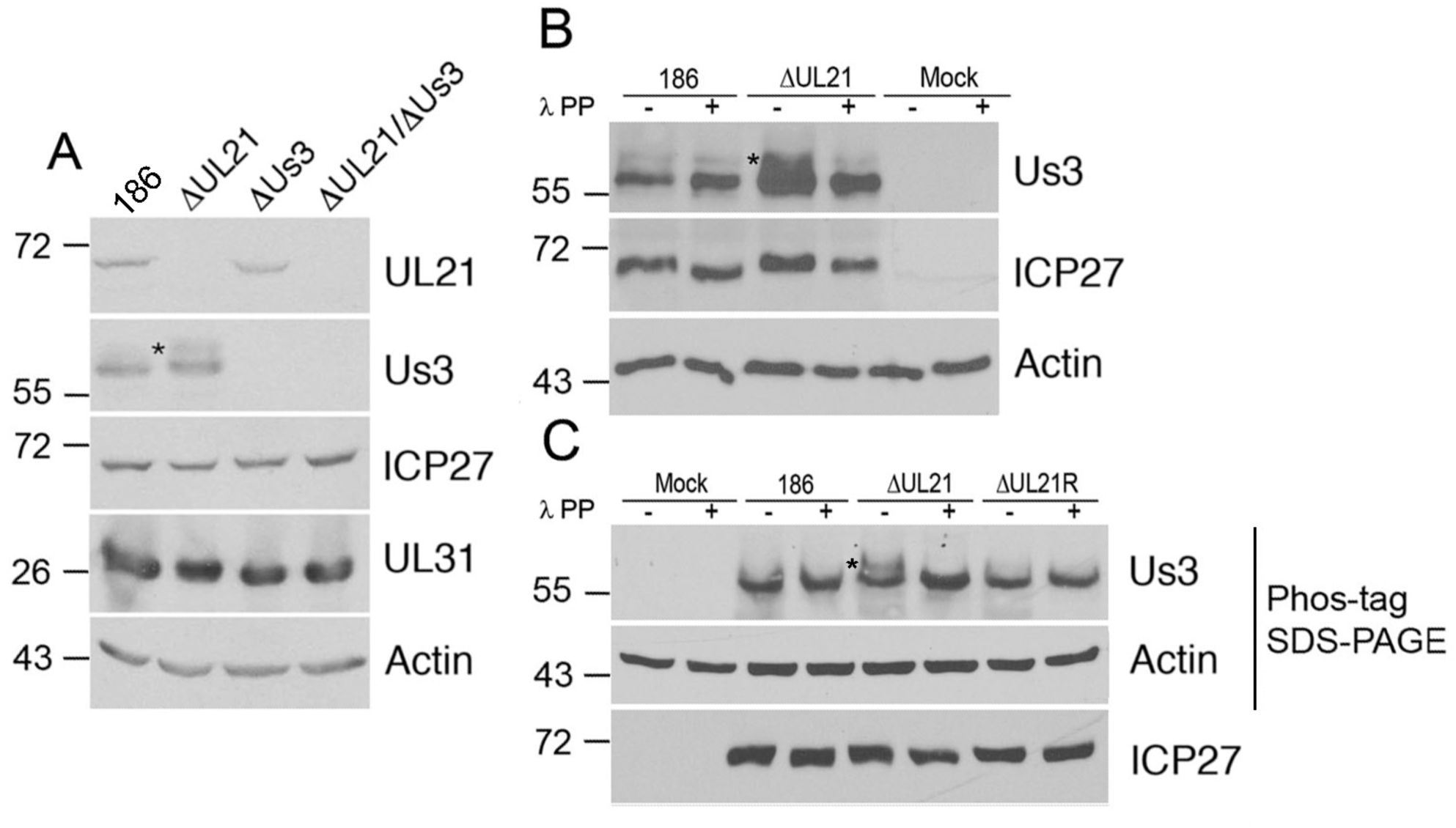
HSV-2 pUs3 kinase is phosphorylated in ΔUL21 infected cells. **(A)** Whole cell lysates prepared at 18 hours post infection from Vero cells infected at an MOI of 3 with the viruses indicated at the top of the panels or from mock infected cells were electrophoresed through polyacrylamide gels and proteins transferred to PVDF membranes. Membranes were probed with antisera indicated on the right side of each panel. Molecular weight markers in kDa are indicated on the left side of the panels. Asterisk indicates a slower migrating species of pUs3 that was prominent in cells infected with ΔUL21. **(B)** Whole cell lysates prepared at 18 hours post infection from Vero cells infected at an MOI of 3 with the viruses indicated at the top of the panels or from mock infected cells were either treated with λ protein phosphatase (λ PP) or untreated, then electrophoresed through polyacrylamide gels and proteins transferred to PVDF membranes. Membranes were probed with antisera indicated on the right side of each panel. Molecular weight markers in kDa are indicated on the left side of the panels. Asterisk indicates a slower migrating species of pUs3 that was prominent in cells infected with ΔUL21 virus. **(C)** Whole cell lysates prepared at 18 hours post infection from Vero cells infected at an MOI of 3 with the viruses indicated at the top of the panels or from mock infected cells were either treated with λ protein phosphatase (λ PP) or untreated, then electrophoresed through a regular (lower panel) or phos-tag SDS polyacrylamide gels and proteins transferred to PVDF membranes. Membranes were probed with antisera indicated on the right side of each panel. Molecular weight markers in kDa are indicated on the left side of the panels. Asterisk indicates a slower migrating species of pUs3 that was prominent in cells infected with ΔUL21 virus.

As phosphorylation of HSV-1 pUs3 was shown to increase its kinase activity [29], we hypothesized that the phosphorylation of pUs3 that occurs in the absence of pUL21 could result in an alteration of pUs3 kinase activity and, consequently, affect the phosphorylation status of pUs3 substrates. To assess alterations in pUs3 kinase activity, Vero cell lysates from 186, ΔUL21, ΔUs3, ΔUL21/ ΔUs3 double mutant, and ΔUL21 and ΔUs3 repair strains (ΔUL21R and ΔUs3R) were analyzed by western blotting using antiserum reactive against protein kinase A (PKA) substrates (Figure 2). The similarities between PKA and pUs3 substrate specificities enables this antiserum to detect pUs3 substrates [25, 30]. Many additional bands were detected by the PKA substrate antiserum in lysates prepared from all virally infected cells in comparison to lysates prepared from mock infected cells, which could be attributed to the action of pUs3 and/or the virally-induced upregulation of PKA activity reported previously [30]. The PKA substrate phosphorylation profiles, in terms of both the number of bands and band intensities, were comparable for all viruses analyzed suggesting that, for the most part, the increased reactivity of the PKA substrate antiserum was due to the action of PKA rather than pUs3. Curiously, we observed a prominent band between 26 and 34 kDa (arrowhead, Figure 2) in ΔUL21 infected cell lysates that was absent in ΔUL21/ ΔUs3 infected cells suggesting that this protein represented a bona fide pUs3 substrate.

**Figure 2.**
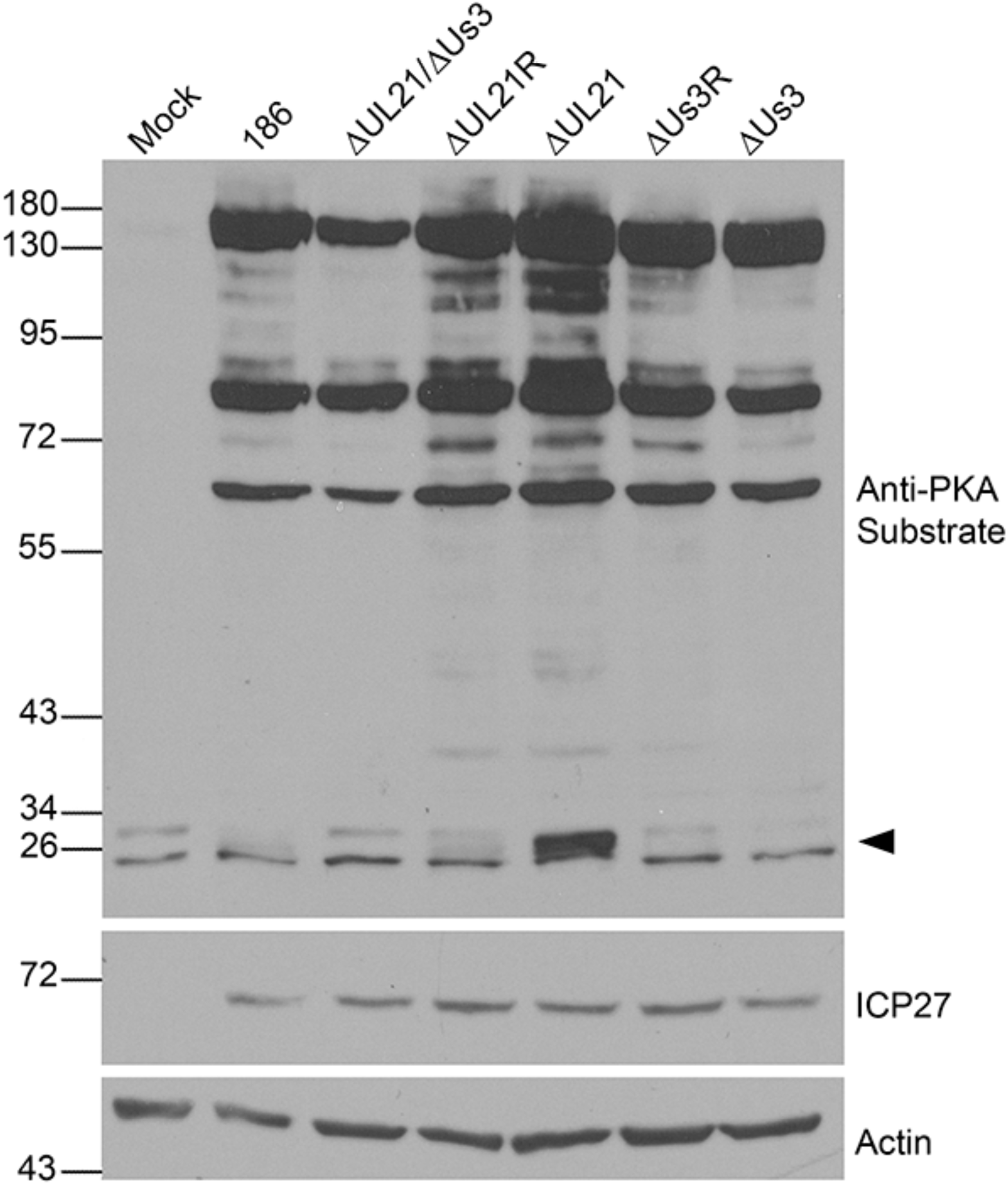
Investigation of phosphorylated PKA/pUs3 substrates in ΔUL21 infected cells. Whole cell lysates prepared at 18 hours post infection from Vero cells infected at an MOI of 3 with the viruses indicated at the top of the panels or from mock infected cells were electrophoresed through polyacrylamide gels and proteins transferred to PVDF membranes. Membranes were probed with antisera indicated on the right side of each panel. Molecular weight markers in kDa are indicated on the left side of the panels. The arrowhead on the right side of the top panel indicates the position of a phosphorylated protein unique to ΔUL21 infected cells.

The migration of the prominent band identified in Figure 2 was consistent with the mobilities of the NEC components pUL31 and pUL34, both of which are known pUs3 substrates [17–20]. Thus, we speculated that this protein could represent either phosphorylated pUL31, phosphorylated pUL34, or both. To test this, 186, ΔUL21 or ΔUL21R infected cell proteins were separated by regular SDS-PAGE and by phos-tag SDS-PAGE and subjected to western blot analysis (Figure 3A). Two distinct phosphorylated forms of pUL34 were resolved by phos-tag SDS-PAGE in all infected cell lysates, both of which were diminished upon phosphatase treatment, and the intensity of the non-phosphorylated species, indicated by the arrowhead in Figure 3A, increased upon phosphatase treatment. We noted that the slowest migrating phosphorylated pUL34 species was more intense in ΔUL21 infected cell lysates in comparison to lysates from 186 and ΔUL21R infected cells. In the case of pUL31, a non-phosphorylated species of pUL31 predominated (arrowhead in Figure 3A) and a single, slower migrating phosphorylated species was resolved in lysates from 186 and ΔUL21R infections. By contrast, three distinct phosphorylated species of pUL31 were readily apparent in lysates prepared from ΔUL21 infected cells. These findings suggest that pUL31 is hyperphosphorylated in ΔUL21 infected cells, as is pUL34, albeit to a lesser extent than pUL31.

**Figure 3.**
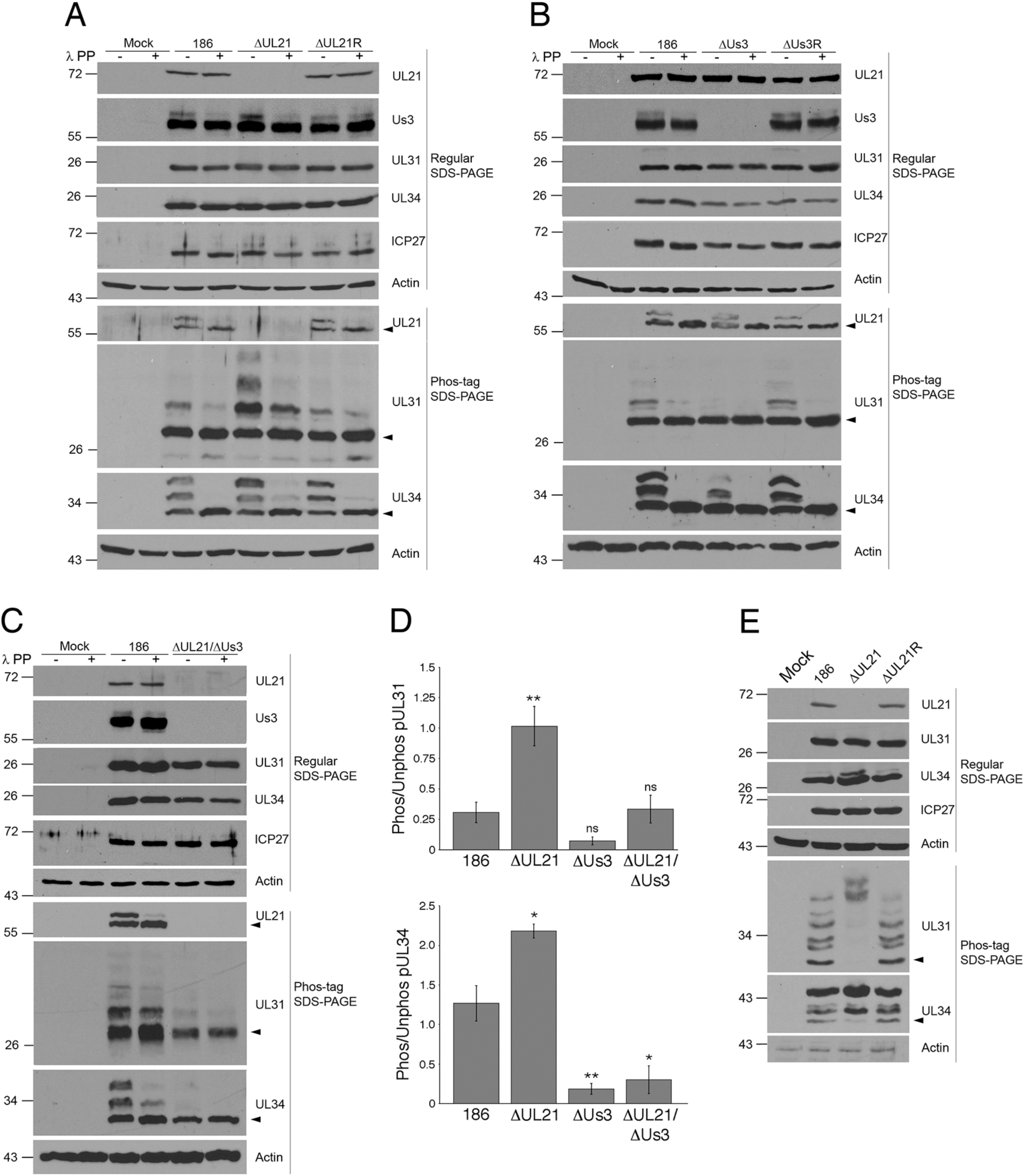
Impact of pUs3 and pUL21 on the phosphorylation of NEC components pUL31 and pUL34. **(A, B and C)** Whole cell lysates prepared at 18 hours post infection from Vero cells infected at an MOI of 3 with the viruses indicated at the top of the panels or from mock infected cells were either treated with λ protein phosphatase (λ PP) or untreated, were electrophoresed through regular or phos-tag SDS polyacrylamide gels and proteins transferred to PVDF membranes. Membranes were probed with antisera indicated to the right of each panel. Molecular weight markers in kDa are indicated on the left side of the panels. Black arrowheads denote the position of unphosphorylated protein species. **(D)** The ratios of phosphorylated to unphosphorylated pUL31 (top graph) or pUL34 (bottom graph) in extracts prepared from Vero cells infected at an MOI of 3 with the viruses indicated on the x-axis was quantitated using Image J software (186 n= 5, ΔUL21 n= 3, ΔUs3 n=3, ΔUL21/ ΔUs3 n=3). Error bars are standard error of the mean. Asterisks indicate significantly different ratios in comparison to the ratios observed in 186 infected cells, determined by Student’s T test; * = P value ≤ 0.05; ** = P value ≤ 0.01; ns = not significant. (**E**) Whole cell lysates prepared at 18 hours post infection from HeLa cells infected at an MOI of 3 with the viruses indicated at the top of the panels or from mock infected cells were electrophoresed through regular or phos-tag SDS polyacrylamide gels and proteins transferred to PVDF membranes. Membranes were probed with antisera indicated to the right of each panel. Molecular weight markers in kDa are indicated on the left side of the panels. Black arrowheads denote the position of unphosphorylated protein species.

HSV-1 pUs3 phosphorylates up to six serines within the N-terminal 50 amino acids of HSV-1 pUL31 [19]. Similarly, the N-terminal 50 amino acids of HSV-2 pUL31 contain seven serines and one threonine located within pUs3 consensus phosphorylation sequences. In keeping with our identification of two distinct phosphorylated forms of pUL34 in HSV-2 infected cells, previous studies identified two residues in HSV-1 pUL34 (threonine 195 and serine 198) that are pUs3 substrates [20], which are both serine residues in HSV-2 pUL34. To test whether the absence of pUs3 would affect the phosphorylation status of the NEC components, lysates from 186, ΔUs3 or ΔUs3R infected cells were subjected to phos-tag SDS-PAGE and western blot analysis (Figure 3B). In agreement with the data shown in Figure 3A, there was a prominent phosphorylated form of pUL31 in 186 and ΔUs3R infected cell lysates. In the absence of pUs3, however, pUL31 phosphorylation was abolished. Furthermore, unlike in 186 and ΔUs3R infected cell lysates, the slowest migrating phosphorylated species of pUL34 was diminished in the absence of pUs3. These findings suggest that while pUs3 is required for pUL31 phosphorylation, it only partially contributes to the phosphorylation of pUL34. To verify that, in the absence of pUL21, pUs3 kinase activity was required for pUL31 and pUL34 hyperphosphorylation, lysates from 186 and ΔUL21/ ΔUs3 double mutant infected cells were subjected to phos-tag SDS-PAGE and western blot analysis (Figure 3C). In the absence of both pUL21 and pUs3, pUL31 hyperphosphorylation was abolished indicating that, as expected, pUs3 was required for pUL31 phosphorylation. Interestingly, no phosphorylated species of pUL34 were observed in the absence of both pUL21 and pUs3 indicating that pUL21 can also influence the phosphorylation state of pUL34 when mediated by kinases other than pUs3. The ratio of phosphorylated pUL31 and pUL34 to their non-phosphorylated species was quantified in cells infected with parental and mutant virus strains (Figure 3D). This analysis confirmed that both pUL31 and pUL34 were significantly hyperphosphorylated in the absence of pUL21.

An unanticipated outcome of these experiments was the observation that roughly half of the pUL21 in cells infected with 186, ΔUs3, ΔUs3R or ΔUL21R was modified by phosphorylation (Figures 3ABC). These findings indicate that pUL21 is phosphorylated by a kinase other than pUs3 and raises the possibility that pUL21 activity is regulated by this post-translational modification.

### Phosphorylation of NEC components by pUs3 influences the nature of NE deformation

The data so far indicated that in the absence of HSV-2 pUL21, NEC components pUL31 and pUL34 were both hyperphosphorylated. We next sought to understand the consequences of pUL31 and pUL34 phosphorylation on NEC function. It is well established that the NEC has an intrinsic ability to deform membranes in the absence of other viral proteins [7, 31, 32]. To investigate the influence of pUs3-mediated phosphorylation of NEC components on nuclear membranes in a reductionist system, we examined the distribution of the NEC in transfected HeLa cells in the presence or absence of pUs3 kinase activity. HeLa cells were chosen for these analyses because of their high transfection efficiency in comparison to Vero cells. As a prelude to these analyses, we verified that HeLa cells infected with pUL21 mutants hyperphosphorylated pUL31 and pUL34 (Figure 3E). These experiments indicated that, similar to what was observed in Vero cells, pUL31 and pUL34 were hyperphosphorylated in the absence of pUL21 in HeLa cells, and arguably, to an even greater extent than was seen in Vero cells.

Previously, we demonstrated that EGFP-pUL31 could be recruited to the nuclear rim of HeLa cells by co-transfecting it with Flag-pUL34 and that, in the case of cells with dim fluorescence, the distribution of the NEC at the nuclear rim was smooth [26]. Cells co-transfected with EGFP-pUL31 and untagged pUL34 showing dim fluorescence had a similar smooth NEC distribution (data not shown). In cells co-transfected with EGFP-pUL31 and either pUL34 or Flag-pUL34 showing bright fluorescence, indicative of robust NEC production, the distribution of the NEC was aberrant with structures that appeared to be located inside of the nucleus (invaginations) (Figures 4B and 5A). Aberrant NEC distribution was also observed in co-transfected cells with robust NEC production in the presence of pUs3, however, the resulting structures appeared to protrude outwards from the nucleus (extravagations) (Figure 4B).

**Figure 4.**
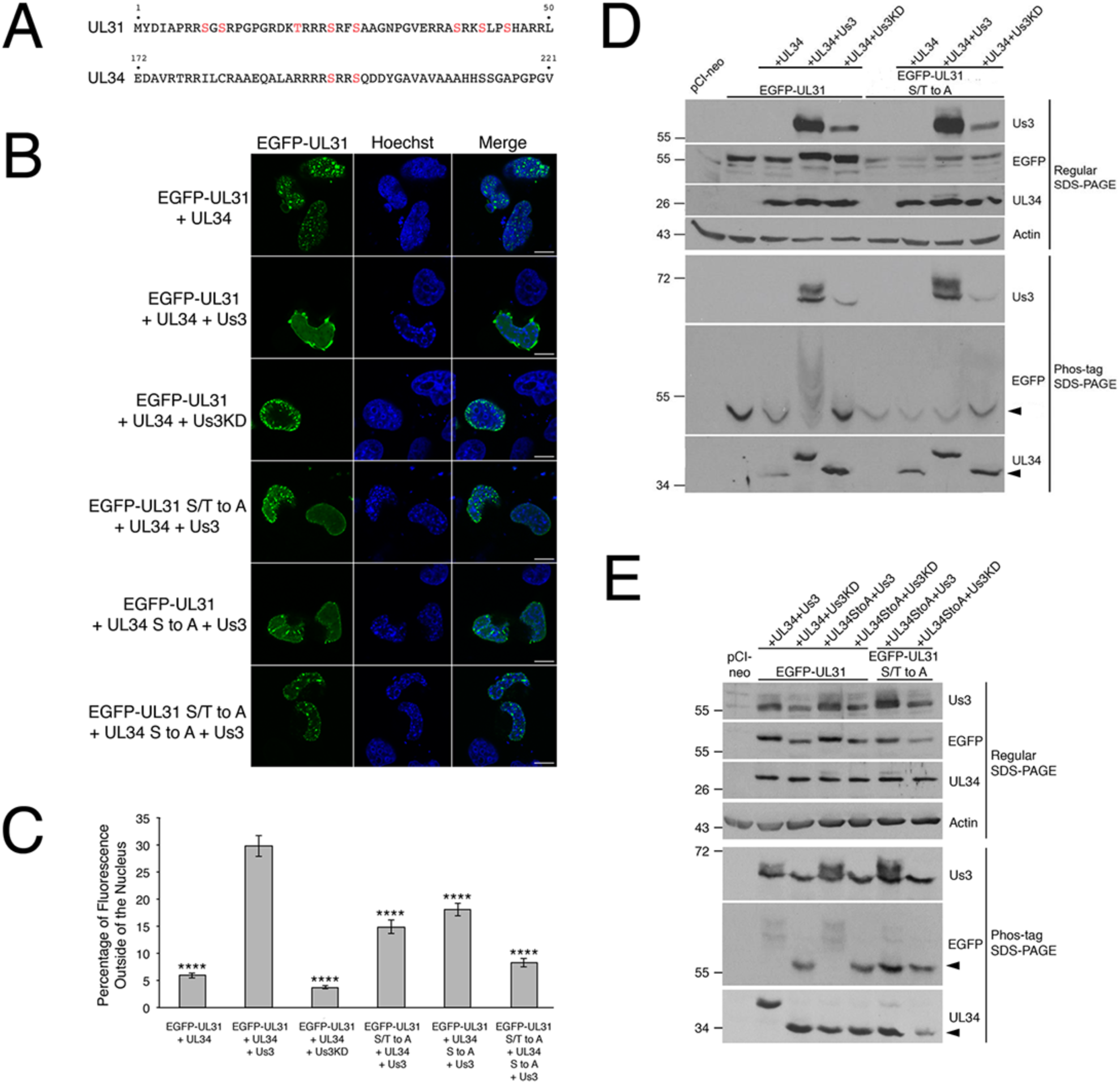
NEC distribution and phosphorylation in transfected HeLa cells. **(A)** Potential pUs3 substrates, indicated in red, reside in HSV-2 pUL31 (top) and in HSV-2 pUL34 (bottom). **(B)** HeLa cells were co-transfected with the expression plasmids indicated on the left of each panel. At 18 hours post transfection, cells were fixed and nuclei were stained with Hoechst 33342. Representative images of cells with bright fluorescence indicative of robust NEC production are shown. Scale bars are 10 μm. **(C)** To quantify the requirements for extravagation formation, HeLa cells were co-transfected with the plasmids indicated on the x-axis, then fixed and stained as described in **(B)**. Images of 70 to 100 cells were captured for each condition and the percentage of fluorescence outside of the nucleus for each image was determined as described in Materials and Methods. Error bars are standard error of the mean. Asterisks denote significant decreases in the percentage of fluorescence outside of the nucleus in comparison to that of co-transfection with EGFP-pUL31, pUL34 and pUs3, determined by Student’s T test; **** = P value < 0.0001. **(D, E)** HeLa cells were transfected or co-transfected with the expression plasmid(s) indicated above each lane. Equal volumes of protein extracts prepared at 18 hours post transfection were electrophoresed through regular or phos-tag SDS polyacrylamide gels and proteins transferred to PVDF membranes. Membranes were probed with antisera indicated on the right of each panel. Molecular weight markers in kDa are indicated on the left side of each panel. Black arrowheads denote the position of unphosphorylated protein species.

**Figure 5.**
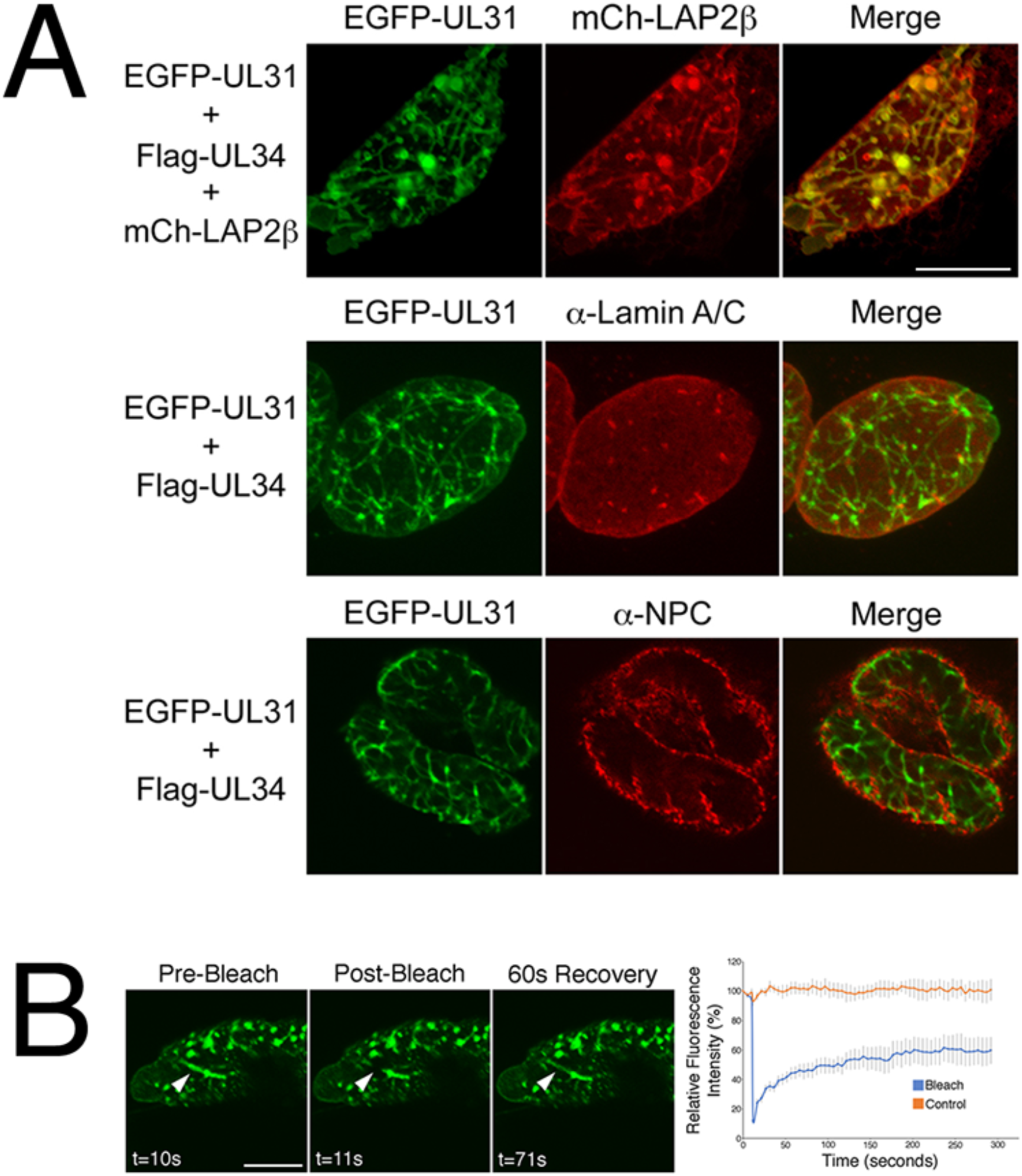
Characterization of invaginations formed in transfected HeLa cells. **(A)** Colocalization analysis. HeLa cells were co-transfected with the expression plasmids indicated on the left of each panel. At 18 hours post transfection, cells were fixed and, in the case of the bottom two panels, stained with the indicated antibodies and Alexa Fluor 568-conjugated secondary antibodies. Representative images of cells with bright fluorescence indicative of robust NEC production are shown. Scale bars are 10 μm. **(B)** Fluorescence recovery after photobleaching analysis. HeLa cells were co-transfected with EGFP-pUL31, pUL34 and pUs3KD. Live cells displaying invaginations were imaged at 18 hours post transfection and selected images of cells immediately before, immediately after, or at a later time after the region indicated by the arrow was photobleached are shown. Scale bar is 10 μm. The corresponding quantification of fluorescence intensity over time of the photobleached region as well as a control region not subjected to photobleaching is shown beside the images. Error bars represent standard error of the mean.

To determine if pUs3 kinase activity was required for switching the NEC distribution in cells from invaginations to extravagations, a mutated version of pUs3 with disrupted kinase activity [25] was co-transfected along with EGFP-pUL31 and pUL34. The resulting NEC distribution pattern resembled that of cells co-transfected with EGFP-pUL31 and pUL34 alone (Figure 4B), indicating that pUs3 kinase activity was required for the switch from invaginations to extravagations. As mentioned above, there are potential pUs3 substrates in the amino terminus of HSV-2 pUL31 and in HSV-2 pUL34 (see Figure 4A). To determine if pUs3-mediated phosphorylation of substrates in pUL31 or pUL34 were required for switching the NEC distribution pattern, a mutated version of EGFP-pUL31 in which serines 9, 11, 24, 27, 39, 42 and 45 and threonine 20 were all changed to alanine (EGFP-pUL31 S/T to A) and a mutated version of UL34 in which serines 195 and 198 were both changed to alanine (pUL34 S to A) were used in co-transfections. In both cases, the resulting NEC distribution pattern more closely resembled that of cells co-transfected with EGFP-pUL31 and pUL34 alone (Figure 4B) than that of cells co-transfected with EGFP-pUL31, pUL34 and pUs3, indicating that pUs3 substrates in pUL31 and pUL34 were also required for the switch from invaginations to extravagations. To quantify the requirements for extravagation formation, the percentage of fluorescence outside of the nucleus in images of co-transfected cells was compared. As shown in Figure 4C, the percentage of fluorescence outside of the nucleus was significantly decreased in co-transfections receiving no pUs3, pUs3KD, EGFP-pUL31 S/T to A and pUL34 S to A in comparison to co-transfections receiving all three wild type proteins, consistent with a requirement for pUs3-mediated phosphorylation of pUL31 and pUL34 in extravagation formation. Extravagations comparable to those observed in cells co-transfected with EGFP-pUL31, pUL34 and pUs3 were not observed in control experiments with cells transfected with pUs3 alone, EGFP-pUL31 alone, pUL34 alone or co-transfected with pUL34 and pUs3 (data not shown). Thus, extravagation formation in transfected cells requires robust production of the NEC, pUs3 kinase activity and serine/threonine substrates in both pUL31 and pUL34.

To determine if there was a correlation between hyperphosphorylation of pUL31 or pUL34 and extravagation formation, protein extracts prepared from transfected HeLa cells were analyzed by phos-tag SDS-PAGE and western blotting. As shown in Figure 4D and Figure 4E, hyperphosphorylation of pUL31 was observed in co-transfected cells that received wild type pUs3. Phosphorylation of pUL34 by pUs3 was also evident in these experiments (see bottom panels of Figure 4D and Figure 4E). In contrast to what was observed in infected cells (Figure 3), only the most slowly migrating form of pUL34 was evident in co-transfected cells. Phosphorylation of pUL31 was specifically lost in co-transfected cells receiving EGFP-pUL31 S/T to A (Figure 4D) and phosphorylation of pUL34 was specifically lost in co-transfected cells receiving pUL34 S to A (Figure 4E).

To determine whether the invaginations observed in cells producing the NEC were comprised of one or both nuclear membranes, we analyzed the colocalization of EGFP-pUL31 with nuclear membrane markers. Good colocalization was observed between EGFP-pUL31 and lamin associated protein 2β, a marker of the INM [33], confirming that invaginations were derived from nuclear membrane(s) (Figure 5A). Fluorescence recovery after photobleaching (FRAP) analysis of nuclear invaginations demonstrated a rapid recovery of EGFP-pUL31 to the photobleached region of the invagination, consistent with nuclear membrane-derived invaginations as opposed to immobile aggregations of protein within the nucleus (Figure 5B). Poor colocalization was observed between EGFP-pUL31 in invaginations and both lamin A/C and nuclear pore complexes (Figure 5A). As invaginations comprised of both ONM and INM would be expected to remain associated with lamins and contain nuclear pore complexes [34], our colocalization analyses indicated that the invaginations formed in transfected cells are comprised of INM only.

To examine the arrangement of nuclear membranes at higher resolution, transmission electron microscopy (TEM) analyses were carried out on transfected cells. In cells transfected with EGFP-pUL31, pUL34 and pUs3KD, invaginations of the INM were observed (Figure 6). These INM invaginations ranged from small herniations at the nuclear periphery (black arrowheads in Figure 6EF) to large herniations containing budded vesicles of uniform size (black arrows in Figure 6DG) to tubular structures extending deep into the nucleus (black asterisks in Figure 6DG) and were not observed in control transfections (Figure 6ABC). In cells co-transfected with EGFP-pUL31, pUL34 and pUs3, extravagations of both ONM and INM (white arrowheads in Figure 7Ai, Aii) or of ONM only (white arrows in Figure 7Aii, Aiii) were observed. In addition to extravagations, long swaths of tightly layered nuclear membranes (zippering) were observed (white asterisks in Figure 7Aiv, Av, Avi). Nuclear membrane extravagations and zippering were not observed in control transfections (Figure 6ABC). FRAP analysis on extravagations formed in transfected cells in the presence of pUs3 also demonstrated a rapid recovery of EGFP-pUL31 to the photobleached region of the extravagation (Figure 7B), in keeping with the membranous nature of the structures that we observed in our TEM analyses of these transfections. Taken together, the results of our transfected cell analyses indicate that phosphorylation of pUL31 and pUL34 by pUs3 influences the nature of nuclear membrane deformation.

**Figure 6.**
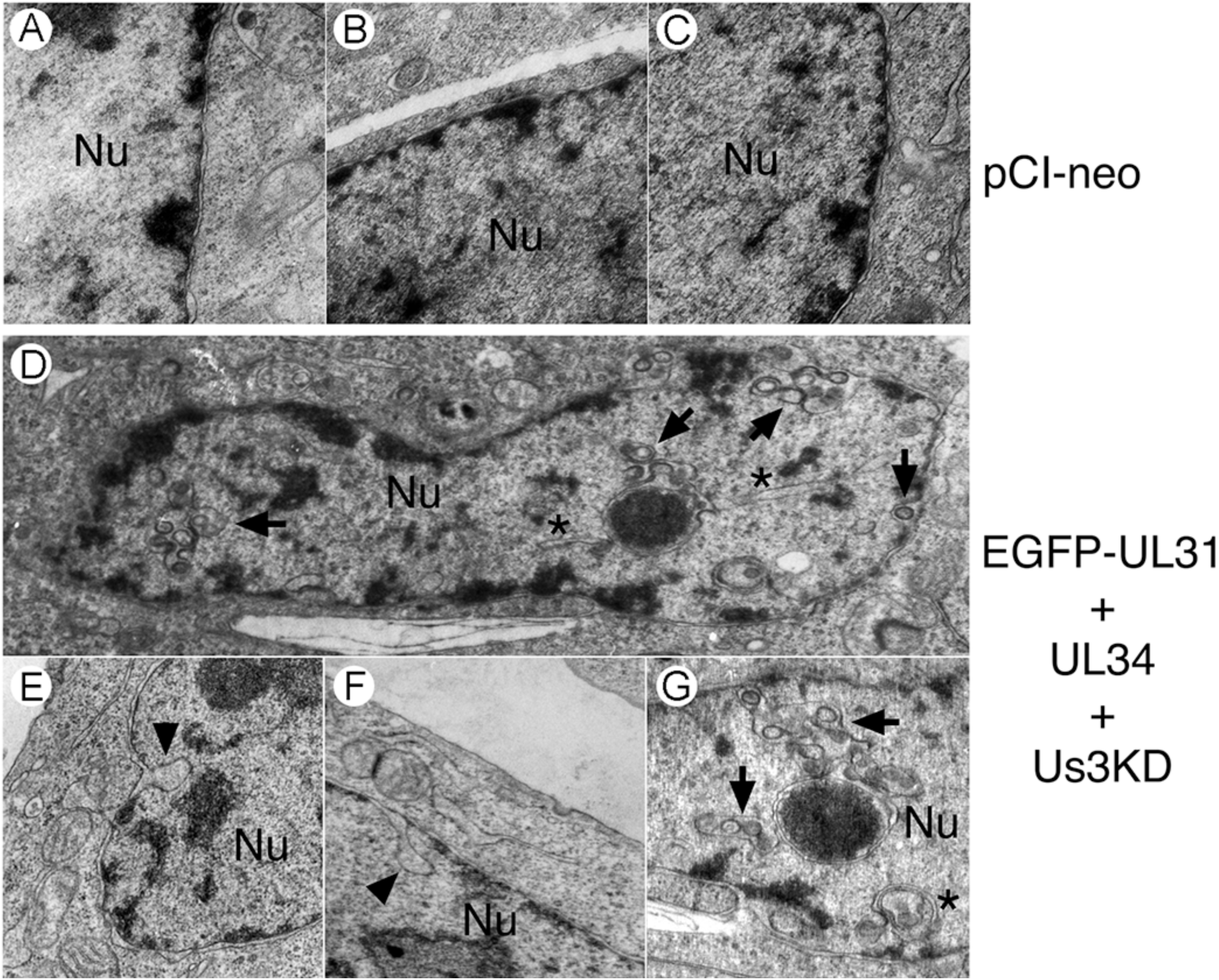
Ultrastructural analysis of invaginations formed in transfected HeLa cells. HeLa cells were transfected with the empty vector, pCI-neo, or with plasmids encoding the indicated proteins. At 18 hours post transfection, cells were fixed and processed for TEM as described in Materials and Methods. HeLa cells co-transfected with plasmids encoding EGFP-pUL31, pUL34 and pUs3KD formed a variety of invaginations that were not observed in control transfections receiving pCI-neo. Black arrowheads indicate invaginations of the inner nuclear membrane (INM), black arrows indicate INM invaginations containing budded vesicles and black asterisks indicate tubular invaginations. Nu indicates the nucleus.

**Figure 7.**
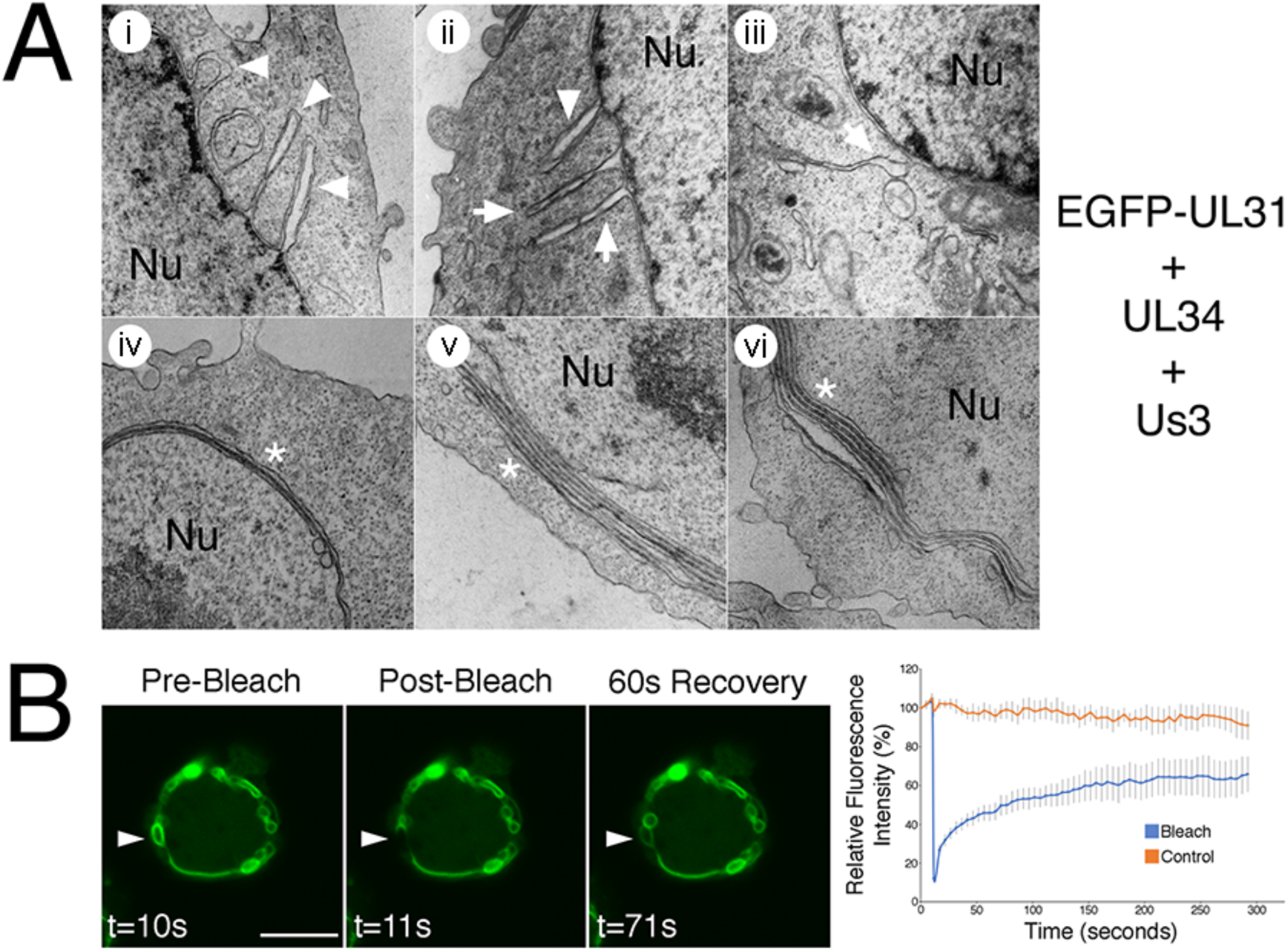
Characterization of extravagations formed in transfected HeLa cells. **(A)** Ultrastructural analysis. HeLa cells were co-transfected with plasmids encoding the indicated proteins. At 18 hours post transfection, cells were fixed and processed for TEM as described in Materials and Methods. HeLa cells co-transfected with plasmids encoding EGFP-pUL31, pUL34 and pUs3 formed extravagations that were not observed in control transfections receiving pCI-neo (see Figure 6). White arrowheads indicate extravagations comprised of both outer nuclear membrane (ONM) and inner nuclear membrane, white arrows indicate extravagations comprised of ONM only and white asterisks indicate zippered nuclear membranes, which were also not observed in control transfections. Nu indicates the nucleus. **(B)** Fluorescence recovery after photobleaching analysis. HeLa cells were co-transfected with EGFP-pUL31, pUL34 and pUs3. Live cells displaying extravagations were imaged at 18 hours post transfection and selected images of cells immediately before, immediately after, or at a later time after the region indicated by the arrow was photobleached are shown. Scale bar is 10 μm. The corresponding quantification of fluorescence intensity over time of the photobleached region as well as a control region not subjected to photobleaching is shown beside the images. Error bars are standard error of the mean.

### pUL21 can regulate pUs3-mediated phosphorylation of NEC components in transfected cells

Our infected cell analyses demonstrated that NEC components were hyperphosphorylated in the absence of pUL21 (Figure 3). To determine if pUL21 was also able to influence pUs3-mediated phosphorylation of NEC components in transfected cells, HeLa cells were co-transfected with EGFP-pUL31, pUL34 and pUs3 in the presence or absence of pUL21 and then the phosphorylation state of NEC components was examined by phos-tag SDS-PAGE. When equal amounts of expression plasmids were used in transfection mixes, very slight decreases in phosphorylation of EGFP-pUL31 and pUL34 were detected in the presence of pUL21 (compare lanes designated high in Figure 8). When the amount of pUs3 expression plasmid included in the transfection mixes was decreased 10-fold (lanes designated low in Figure 8), a much more obvious decrease in phosphorylation of EGFP-pUL31 was detected specifically in the presence of pUL21. The decrease in phosphorylation of pUL34 in the presence of pUL21 was comparable to that observed in the absence of pUL21 when lower amounts of pUs3 were used in co-transfections. These results indicate that pUL21 is able to regulate pUs3-mediated phosphorylation of NEC components in the absence of additional viral proteins.

**Figure 8.**
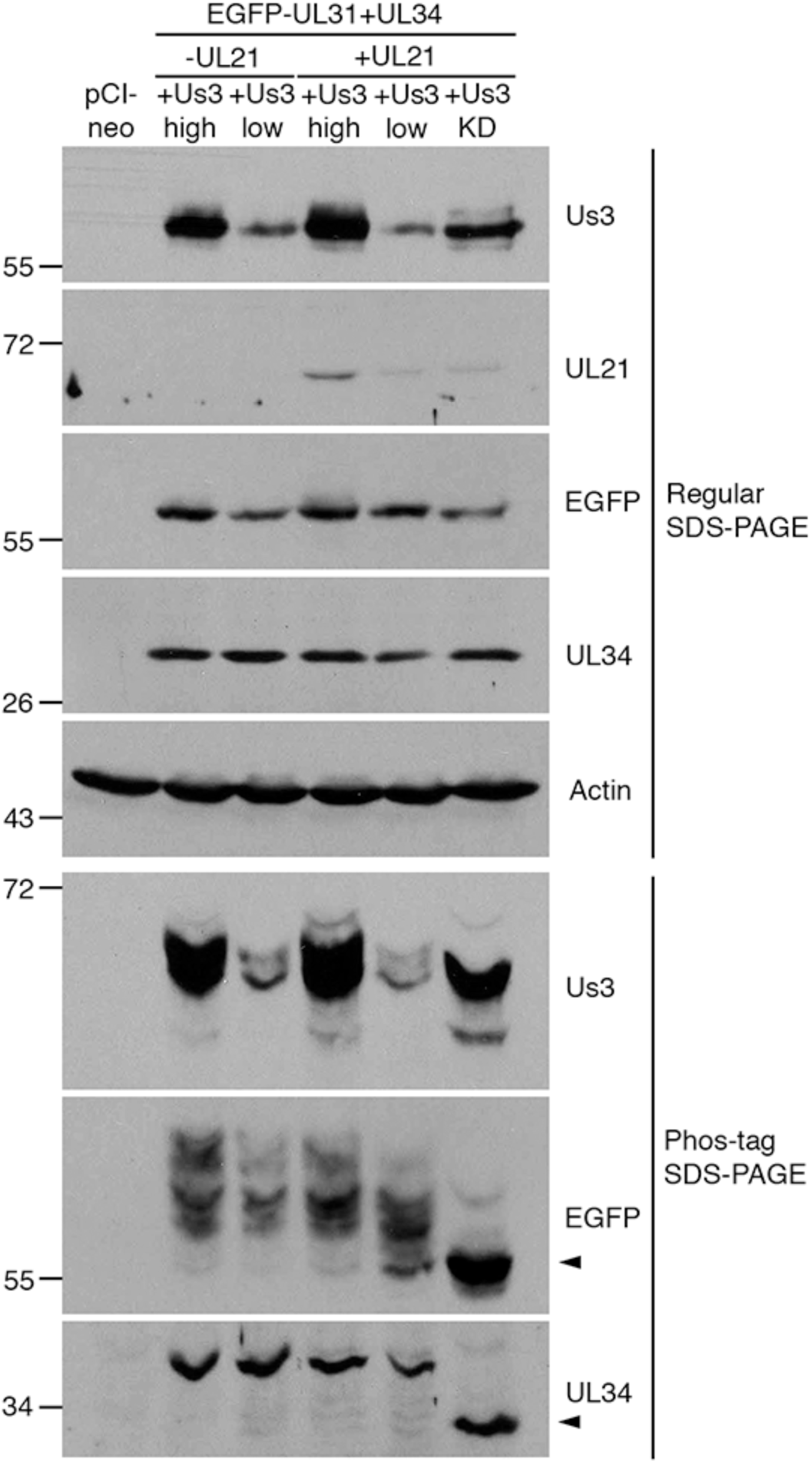
Impact of pUL21 on pUs3-mediated phosphorylation of NEC components in transfected cells. HeLa cells were transfected or co-transfected with the expression plasmid(s) indicated above each lane. In the case of samples designated high, equal amounts of each expression plasmid were included in transfection mixes. In the case of samples designated low, 10-fold less pUs3 expression plasmid was included in transfection mixes and an appropriate amount empty pCI-neo vector was included to normalize the amount of DNA in the transfection mix. In transfection mixes lacking pUL21 expression plasmid, an equivalent amount of empty pCI-neo vector was substituted to normalize the amount of DNA in the transfection mix. Equal volumes of protein extracts prepared at 18 hours post transfection were electrophoresed through regular or phos-tag SDS polyacrylamide gels and proteins transferred to PVDF membranes. Membranes were probed with antisera indicated on the right of each panel. Molecular weight markers in kDa are indicated on the left side of each panel. Black arrowheads denote the position of unphosphorylated protein species.

## Discussion

The NEC functions to envelope C-capsids at the INM. This NEC activity must be regulated in herpesvirus infected cells to prevent excessive INM vesiculation and invagination at sites where C-capsids are absent. In this study, we investigated the roles of HSV-2 pUL21 and pUs3 in the regulation of the NEC and the consequences of pUL31 and pUL34 phosphorylation in remodeling the NE. In the absence of pUL21, pUs3 was hyperphosphorylated (Figure 1) prompting us to investigate the phosphorylation status of pUs3 substrates in cells infected with a pUL21 mutant HSV-2 strain, ΔUL21 (Figure 2). Knowing that: 1) ΔUL21 has a nuclear egress deficiency [22]; the NEC is mislocalized in ΔUL21 infected cells [16]; and 3) NE disruptions observed in ΔUL21 infected cells required pUL31 [16], we examined pUs3-dependent phosphorylation of pUL31 and pUL34 in the absence of pUL21 (Figure 3).

The HSV serine/threonine kinase, pUs3, performs several critical functions for the virus, from blocking apoptosis to regulation of nuclear egress [11]. In PRV [15], equine herpesvirus 1 [12], Marek’s disease virus [14], HSV-1 [13] and HSV-2 [16] Us3 mutants, INM invaginations containing accumulations of PEVs were observed. Baines and colleagues studying the HSV-1 pUs3/pUL31 catalytic relationship, noted that mutation of pUL31 N-terminal serine residues that are pUs3 substrates had a profound influence on NEC activity. When these residues were mutated to alanine, the phenotype resembled that of Us3 null viruses with PEVs accumulating within herniations of the INM. Conversely, phosphomimetic substitution of these serine residues to glutamic acid prevented C-capsid envelopment at the INM. These findings suggest that the hypophosphorylation of pUL31 enables NEC activation and capsid envelopment [19]. Our results are consistent with these findings and extend our understanding of nuclear egress to include a role for pUL21 in the negative regulation of the NEC by modulating the phosphorylation status of pUs3, pUL31 and pUL34.

How might pUL21 regulate the phosphorylation of pUs3, pUL31 and pUL34? Based on our observations that pUs3 is hyperphosphorylated in the absence of pUL21 and that this was accompanied by hyperphosphorylation of pUs3 substrates, pUL31 and pUL34, a plausible explanation is that pUs3 hyperphosphorylation leads to increased pUs3 activity towards pUL31 and pUL34 (Figure 9). While we have not established a relationship between pUs3 hyperphosphorylation and increased kinase activity, it is known that autophosphorylation of HSV-1 pUs3 at serine 147 (S147) enhances pUs3 kinase activity [27]. HSV-2 pUs3 contains an aspartic acid at position 147 (D147) and it is presently unclear which HSV-2 pUs3 residue is hyperphosphorylated in the absence of pUL21. Intriguingly, mutation of D147 in HSV-2 pUs3 inhibits kinase activity [35]. Kawaguchi and colleagues suggested that HSV-1 pUs3 phosphorylation of S147 allows for a reversible regulation of kinase activity, while HSV-2 pUs3 D147 mimics the constitutively active state of the enzyme and is required for maintenance of kinase activity [35]. While HSV-2 pUs3 is phosphorylated in transfected cells, a kinase-dead version of pUs3 is not, raising the possibility that HSV-2 pUs3 phosphorylates itself. However, this interpretation is confounded by the lack of pUs3 consensus phosphorylation sites within HSV-2 pUs3, so it is possible that pUs3 kinase activity is required to enable a cellular kinase to phosphorylate HSV-2 pUs3. The identification of the amino acid in HSV-2 pUs3 that is phosphorylated and how this modification influences pUs3 activity is an outstanding question that, when answered, will advance our understanding of HSV-2 pUs3 regulation and function.

**Figure 9.**
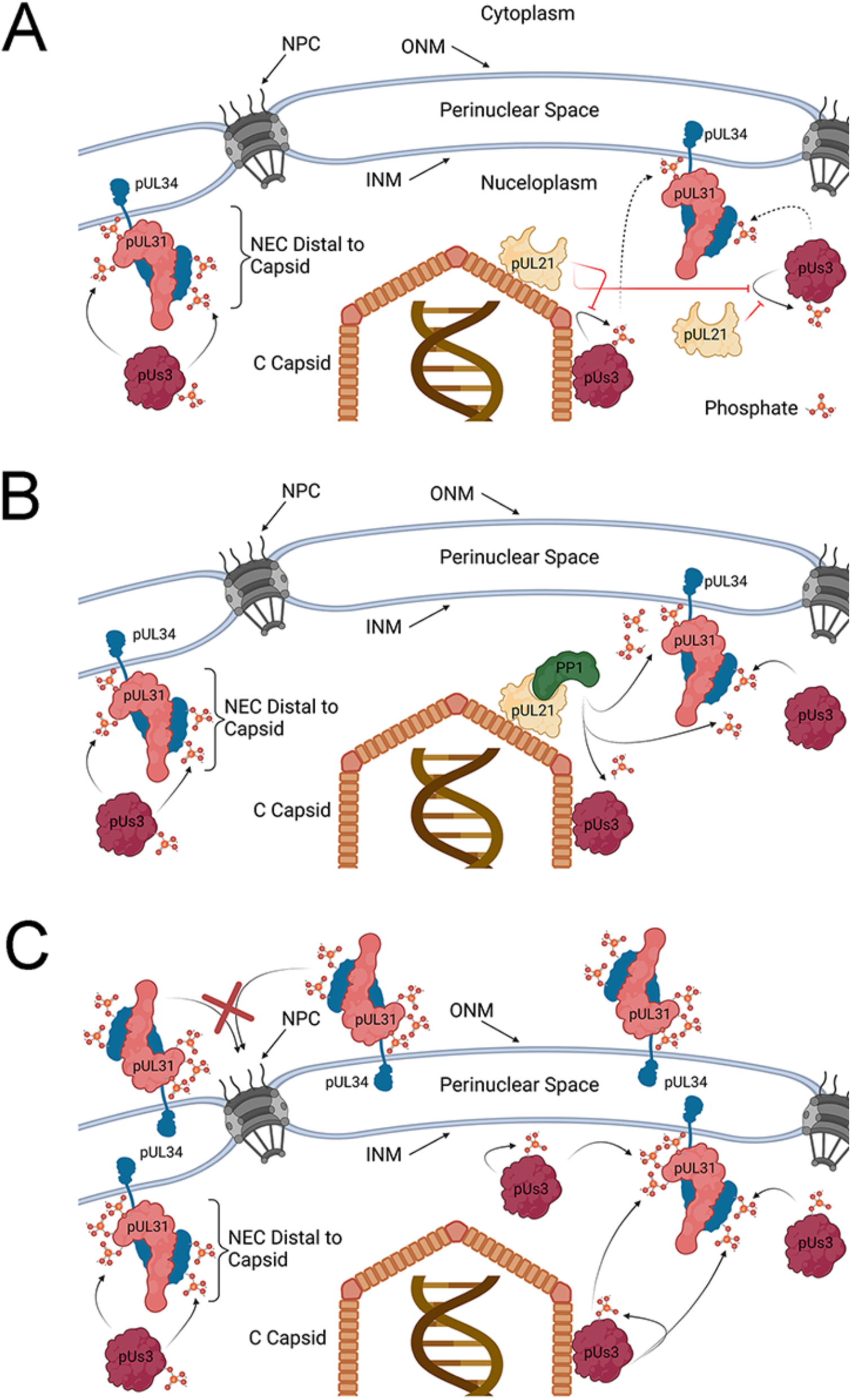
Models for regulation of HSV-2 NEC phosphorylation by pUL21. **(A)** C-capsid associated pUL21 prevents the autophosphorylation of pUs3 (red lines) leading to reduced pUs3 kinase activity (dashed arrows), low level phosphorylation of capsid-proximal pUL31 and pUL34 and corresponding enhancement of NEC primary envelopment and INM invagination activity. Capsid distal NECs are not subject to pUL21 inhibition of pUs3 autophosphorylation and kinase activation, and so, these NECs are more heavily phosphorylated and lack NEC envelopment activity. **(B)** C-capsid associated pUL21 in complex with PP1 facilitate the dephosphorylation of capsid-proximal pUL31, pUL34 and pUs3 leading to enhancement of NEC primary envelopment and INM invagination activity. Capsid distal NEC components are not subject to pUL21/PP1 mediated dephosphorylation and thus these NECs are more heavily phosphorylated and lack NEC envelopment activity. Models A and B are not mutually exclusive and may operate in concert. **(C)** In the absence of pUL21, pUs3 kinase activity is unrestricted and dephosphorylation of pUL31, pUL34 and pUs3 is inefficient leading to hyperphosphorylation of the NEC and inhibition of NEC primary envelopment activity. Hyperphosphorylation of pUL31 interferes with the nuclear import of pUL31/pUL34 NEC complexes in the ONM (red X). Resulting accumulation of hyperphosphorylated NECs in the ONM may lead to nuclear envelope extravagations seen in the absence of pUL21. Outer nuclear membrane (ONM), inner nuclear membrane (INM), nuclear pore complex (NPC). Figure created with BioRender.com

An alternative explanation for the hyperphosphorylation of pUs3, pUL31 and pUL34 in ΔUL21 infected cells is that pUL21 may promote their dephosphorylation (Figure 9B). This idea is strongly supported by a recent unpublished study from Graham and co-workers that demonstrated HSV-1 pUL21 is an adaptor protein for protein phosphatase 1 (PP1) and furthermore that pUL21 has a penchant for delivery of PP1 to pUs3 substrates to mediate their dephosphorlyation [36]. The fact that pUL21 and pUs3 are only found in members of the *Alphaherpesvirinae* raises the intriguing possibility that pUL21 exists, in part, to specifically counter the activities of pUs3. Comparisons of pUs3 substrates with substrates targeted by pUL21/PP1 complexes should provide a clearer picture of the scope and implications of these opposing activities. pUL21-mediated dephosphorylation of pUs3, pUL31 and pUL34 is consistent with our findings demonstrating hyperphosphorylation of these molecules in ΔUL21 infected cells (Figure 2 and the reversal of pUL31 phosphorylation observed in the presence of pUs3 and pUL21 in transfected cells (Figure 8). In the course of these studies we noted that pUL21 is phosphorylated in infected cells by a kinase other than pUs3 (Figure 3). The pUL21 residue that is phosphorylated has yet to be identified, but it is tempting to speculate that this modification may regulate pUL21 activities, subcellular localization and/or interactions with other molecules.

The data presented herein suggest that the function of NEC components is tightly and dynamically regulated by pUs3 and pUL21. Our findings outlined in Figure 3 demonstrate that, in wild type infected cells, pUL31 exists as a mixed population of mostly unphosphorylated and lightly phosphorylated species. pUL34 also exists as a mixture of unphosphorylated and phosphorylated forms. This diverse, moderately phosphorylated population of NEC components underlies the smooth distribution, and the largely quiescent activity, of the NEC at the nuclear rim of wild type HSV-2 infected cells. It may be that capsid-associated pUL21 in complex with an egressing C-capsid locally counteracts pUs3 activity to promote hypophosphorylation and corresponding activation of NEC components, creating a capsid-proximal environment conducive for primary envelopment (Figure 9AB). In the absence of pUs3, as observed in ΔUs3 infected cells, phosphorylation levels of the NEC cannot be maintained leading to NEC hyperactivity manifested by invaginations of the INM, robust primary envelopment and accumulations of PEVs distal to the ONM. This NEC activity could be recapitulated in transfected cells where INM invagination and vesiculation were observed under conditions where pUL31 and/or pUL34 were hypophosphorylated (Figures 4, 5 and 6).

Significant increases in NE extravagations into the cytoplasm were observed when pUL31 and pUL34 expression plasmids were co-transfected with a plasmid expressing pUs3 (Figures 4 and 7). These NE extravagations correlated with enhanced phosphorylation of pUL31 and pUL34 (Figure 4) and were consistent with the structures formed in cells infected with pUL21 mutants derived from multiple HSV-2 and HSV-1 strains [16]. It is possible that an increase in the number and the abundance of the NEC phosphoforms in ΔUL21 infected cells underlies an alteration in NEC activity precluding primary envelopment of capsids and inducing NE extravagations (Figure 9C). One possible explanation for the emergence of the NE extravagations is that pUL31/pUL34 accumulation at the ONM causes the deformation of the ONM through mechanisms analogous to hypophosphorylated NEC components at the INM. The N-terminal region of pUL31, that also contains serine and threonine residues phosphorylated by pUs3, contains a lysine and arginine rich nuclear localization signal (NLS) [3, 26].

Phosphorylation of residues in the pUL31 N-terminus would be expected to interfere with the positively charged NLS. Indeed, HSV-2 pUL31 mutants bearing glutamic acid substitutions of the N-terminal serines and threonine are not efficiently imported into the nucleus (data not shown). Therefore, pUs3 dependent hyperphosphorylation of the pUL31 N-terminus in ΔUL21 infected cells could preclude its proper localization and/or re-location to the INM from the ONM/ER.

In summary, we have established pUL31 and pUL34 as substrates of HSV-2 pUs3 and demonstrated that differential pUL31 and pUL34 phosphorylation underlies different modifications of the NE, using both viral infection and a reductionist co-transfection system. NEC hyperphosphorylation in the absence of pUL21 interferes with NEC activities that mediate the primary envelopment of capsids at the INM and provides an explanation for why HSV-2 strains lacking pUL21 are defective in nuclear egress. A thorough understanding of how NEC activity is controlled during infection may yield strategies to disrupt this fundamental step in the herpesvirus lifecycle, providing the basis for novel antiviral therapies.

## Acknowledgments

This work was supported by the Canadian Institutes of Health Research operating grant 407982, Natural Sciences and Engineering Research Council of Canada Discovery Grant RGPIN-2018-04249 and Canada Foundation for Innovation award 16389 to BWB. JHM was supported in part by a Franklin Bracken Fellowship awarded by Queen’s University. MAG was supported in part by a Natural Sciences and Engineering Research Council of Canada undergraduate summer research award. The funders had no role in study design, data collection and interpretation, or the decision to submit the work for publication. We thank Dr. Tokuko Haraguchi (Osaka University) for providing plasmid encoding EGFP-LAP2β, Dr. Xiaohu Yan (Queen’s University) for assistance with preparing samples for TEM analyses, Dr. Fanrui Meng (Olympus Canada) for assistance with extravagation quantification, Trinity Vey for help with quantitation of pUL34 and pUL31 phosphorylation and Kristin Piche for technical assistance with plasmid construction. We are grateful to Dr. Stephen Graham and members of the Banfield laboratory for helpful comments on the manuscript.

